# Basal-like and Classical cells coexistence in pancreatic cancer revealed by single cell analysis

**DOI:** 10.1101/2020.01.07.897454

**Authors:** Natalia Juiz, Abdessamad Elkaoutari, Martin Bigonnet, Odile Gayet, Julie Roques, Rémy Nicolle, Juan Iovanna, Nelson Dusetti

## Abstract

Pancreatic ductal adenocarcinoma (PDAC) is composed of stromal, immune and epithelial cells. Transcriptomic analysis of the epithelial compartment allows a binary classification into mainly two phenotypic subtypes, classical and basal-like. However, little is known about the intra-tumor heterogeneity of the epithelial component. Growing evidences suggest that this two side phenotypic segregation is not so clear and that both could coexist in a single tumor. In order to elucidate this hypothesis, we performed single-cell transcriptomic analyses using combinational barcoding on epithelial cells from 6 different classical PDAC obtained by Endoscopic Ultrasound (EUS) with Fine Needle Aspiration (FNA). In order to purify the epithelial compartment, PDAC were grown as Biopsy Derived Pancreatic Cancer Organoids. Single cell transcriptomic analysis allowed the identification of 4 main cell clusters present in different proportions in all tumors. Remarkably, although these tumors were classified as Classical, one of the clusters corresponded to a basal-like. These results depict the unanticipated high heterogeneity of pancreatic cancers and demonstrated that basal-like cells with a high aggressive phenotype are more widespread than expected.

## Introduction

With a survival rate of 5 years in less than 8% of the cases (Siegel et al., 2018) pancreatic ductal adenocarcinoma (PDAC) is still one of the most lethal cancers. A principal problem facing this disease is its heterogeneity that results as a consequence of the combination of genetic, epigenetic, and micro-environmental factors (Lomberk et al., 2019, 2018; Yachida and Iacobuzio-Donahue, 2013). Recently, two main PDAC subtypes have been identified by molecular characterization: 1-classical, that are more frequently resectables, presenting a higher level of differentiation, often associated with fibrosis and inflammation; and 2-basal-like, with a poorest clinical outcome and a loss of differentiation (Moffitt et al., 2015; Nicolle et al., 2017). This well-established binary classification could be controverted if cells from a unique tumor contain both phenotypes at the same time. In fact, in addition to the tumor differences between patients that argues in favor of stratification for personalizing PDAC treatments, it is also absolutely necessary to consider the intra-tumor differences since they are playing key roles in the evolution of tumors (i.e.: they can conduce to clonal selection of resistant cells and to the relapse so frequently observed after the first line of chemotherapy).

Single-cell analysis by transcriptomics is nowadays a powerful strategy to determine the intra-tumor heterogeneity, however we need to bypass two main challenging difficulties: The first one is to obtain pure epithelial transformed cells. To do that, we specifically amplified these cells by a few passages in three dimensional (3D) *ex vivo* culture (Tiriac et al., 2019). Three dimensional (3D) cultures of PDAC as tumoral organoids preserve and allow the amplification only of epithelial cancerous cells and create complex structures with polarized cells that recapitulate tumor morphology and allow the communication among different cells within each microtumor (Boj et al., 2015; Tuveson and Clevers, 2019). In order to avoid excessive cell culturing, organoids were directly obtained from primary PDAC samples by endoscopic fine-needle aspiration (EUS-FNA). The second difficulty is to avoid inducing transcriptional modifications in the samples to be studied during library preparation. Most scRNA-seq methods require the capture of viable single cell by cell sorters (Picelli et al., 2013), droplet-based microfluidics (Klein et al., 2015; Macosko et al., 2015) or microwells. These manipulations can completely alter the transcriptional shape of cells, for this reason we decided to use a combinatorial barcoding method, known as the SPLIT-seq approach, that do not require cell physical isolation or complex and long manipulations and cells can be immediately fixed after dissociation (Cao et al., 2019, 2017; Rosenberg et al., 2018). The combinational barcoding also presents an additional advantage regarding its compatibility with big longitudinal sample collections because of its reduced batch effects.

Therefore, in this work, we performed single cell analysis by the SPLIT-seq technology on Biopsy Derived Pancreatic Cancer Organoids (BDPCO) to unravel intratumoral heterogeneity exclusively in the epithelial compartment of 6 PDAC patients.

## Results

### Phenotype characterization of organoids

Six consecutive BDPCOs were prepared and characterized by histologic, immunostaining and transcriptomic analysis after a maximum of 4 *in vitro* passages (Figure 1). In all cases hematoxylin and eosin stained organoids exhibited the formation of glandular architectures with lumen and mucus secretion in all samples. Cells observed were polarized based on immunofluorescent staining for type IV collagen (COL IV) and zona occludes (ZO-1), markers of basement and apical membranes, respectively (Figure 1A and 1B). These anatomophatological characteristics suggests that all cells in the organoids come from the epithelial compartment and are organized as well differentiated glands suggesting that they belong phenotypically to the classical PDAC subtype. In order to confirm their transcriptomic phenotype, a profiling was performed using bulk RNA-seq on these 6 BDPCOs. A 50 gene molecular signature able to identify classical tumors was defined based on the recently Basal-like/classical classification proposed by Nicolle et al, 2017 (Nicolle et al., 2017). Accordingly, to the histologic features, the transcriptomic analysis shows that all BDCPOs present high expression of transcripts associated to the classical subtype indicating that they belong and preserve the classical PDAC phenotype (Figure 1C).

**Figure 1.**
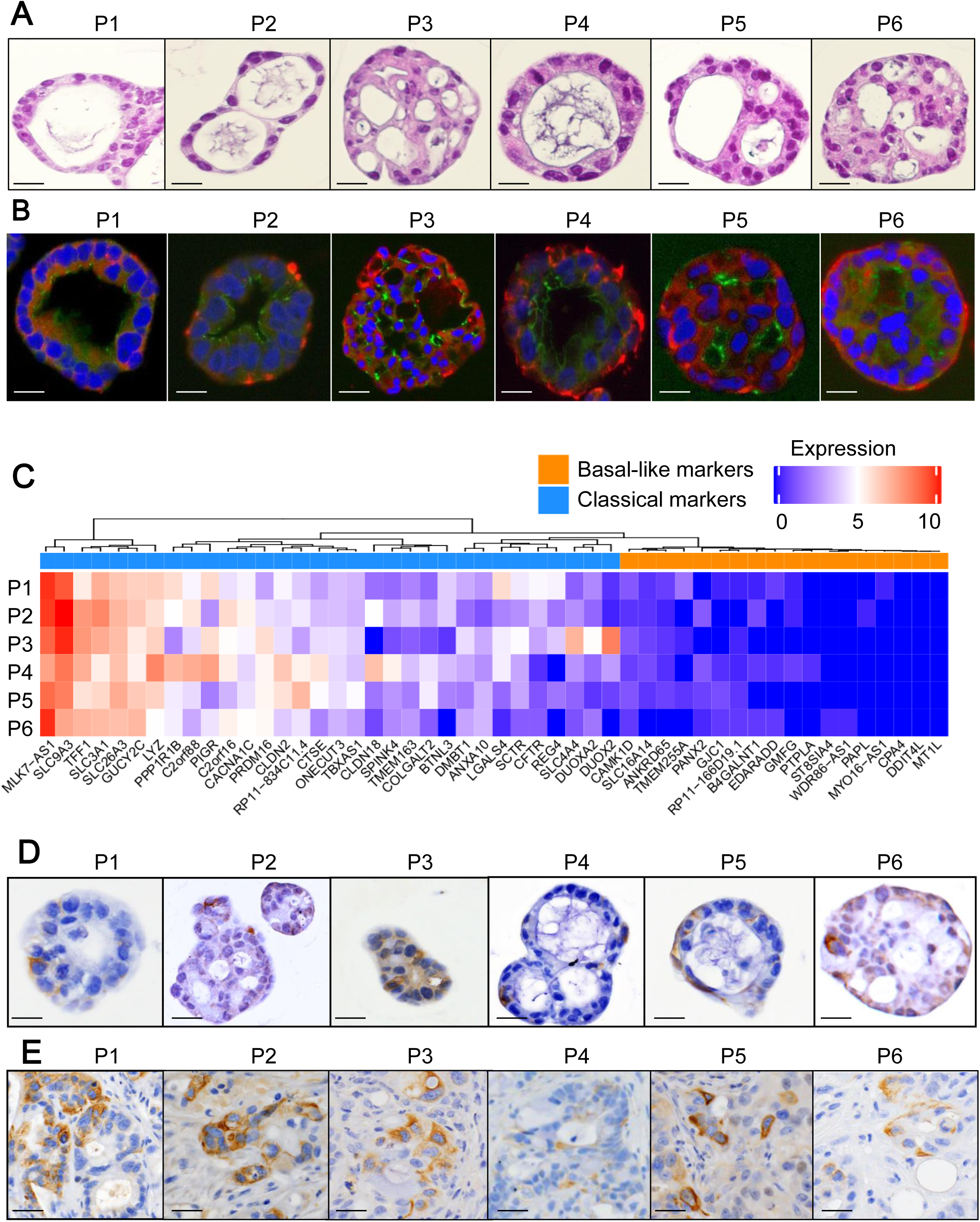
Characterization of PDAC-derived organoids. **A.** Histological characterization of the 6 BDPCO by H&E staining. **B.** Organoids show the presence of glandular structures composed by an apical pole (marked by ZO-1 in green) and a basolateral membrane (marked by COL-IV in red). Scale bar is 50 µm. **C.** Heatmap showing the expression of a 50 gene molecular signature able to identify classical tumors in all six patients. **D.** Immunohistochemical characterization of the six BDPCO with anti-vimentin antibodies. **E.** Immunohistochemical characterization of the six BDPCO derived PDX with anti-vimentin antibodies.

Then, in order to study PDAC heterogeneity, we started by characterizing vimentin (VIM) expression on these BDPOs by immunohistochemistry. We observed that most of the organoids that we obtained directly from patients are clearly heterogeneous presenting concomitantly VIM+ and VIM-cells (Figure 1D). Then, we confirmed that this heterogeneity is also present when organoids were grown *in vivo* as Patient Derived Xenografts (PDX). To do this, we injected these organoids in nude mice and found that this heterogeneous expression of VIM was conserved for at least two passages in PDX (Figure 1E). As VIM is a good basal-like marker, we hypothesized that heterogeneous organoids could contain concomitantly basal-like and classical cells indicating that stratifying tumors in a binary classification (basal-like or classical) could not be as exact as previously supposed.

### Setting a performant scRNA-seq analysis by combinational indexing

In order to perform single cell analysis, organoids cultures from 6 patients were dissociated in a single-cell suspension with a slight protease treatment (see M&M). Two thousands cells from each patient (12,000 total cells) were analyzed by single-cell combinational indexing using the SPLIT-seq technology as previously described (Rosenberg et al., 2018). Cells were formaldehyde-fixed and frozen immediately after treatment. Cultures were expanded to 8 wells on a 48 well/plate and the first indexes were added by retro-transcription. This step allowed the identification of each cell origin knowing that the first barcodes were sample-specific. This first barcoding round was followed by two consecutive ligation steps of barcodes in two 96 well/plates, resulting in a total of 442,368 (48 × 96 × 96) different barcode combinations (Figure 2A). Following the library construction and sequencing, we obtained an excellent performance (in a single cell context) of uniquely mapped reads representing 64.67% of the total reads (982,396,428). After filtering at 15,000 reads by cell, 8,934 individual cells were validated to be considered for analysis with a median of 27,220 reads per cell. Most of the cells (74.45% from the total) were included indicating the high efficiency and quality of barcoding obtained with the modified SPLIT-seq method setup for this work. The *Pearson* correlation between read counts and genes was next to 1 (0.94) and the number of detected genes and reads were similarly distributed across patients. Both parameters indicate a good and unbiased library preparation and amplification (Figure S1A and B).

**Figure 2.**
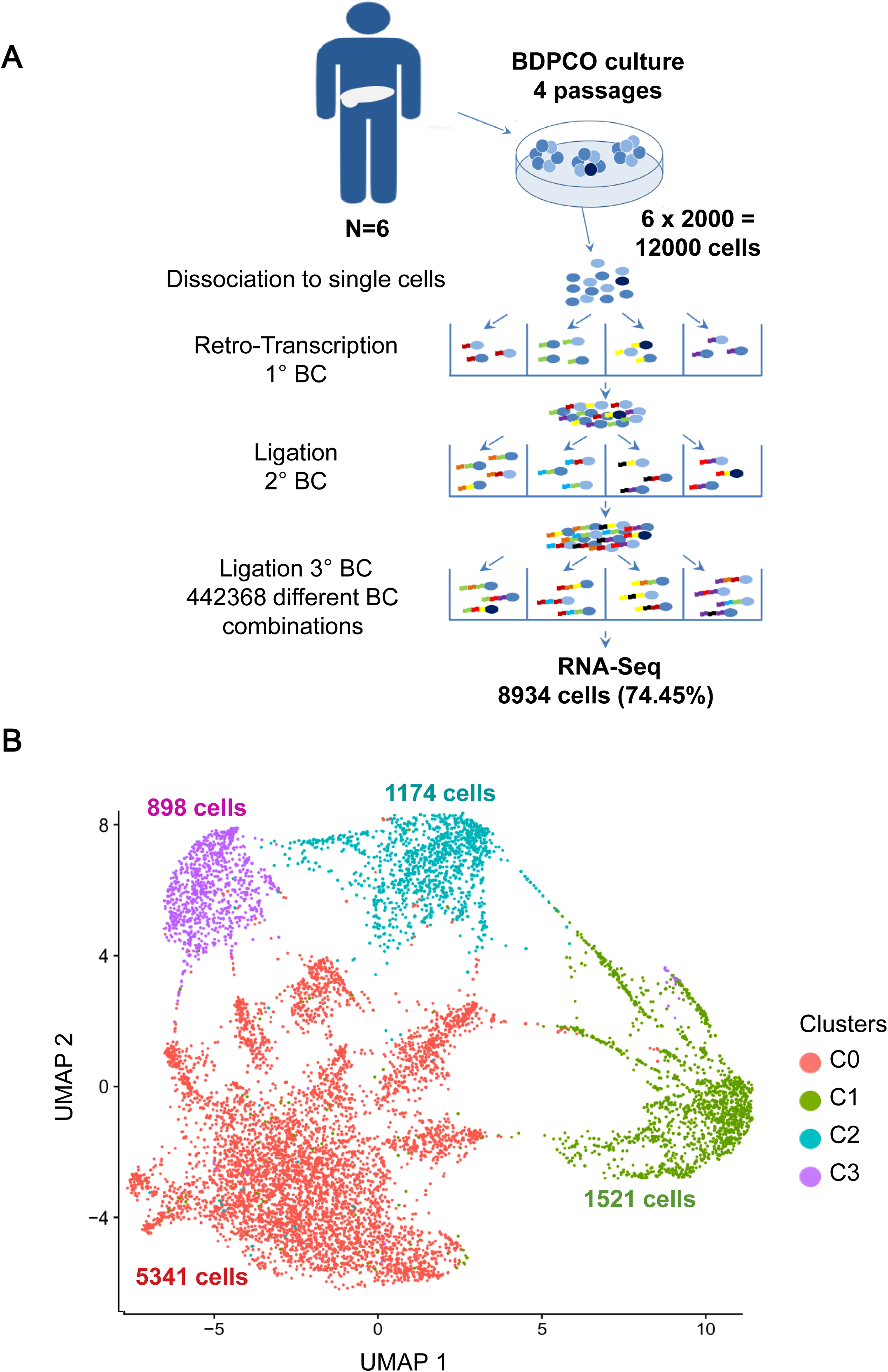
Identification of cell clusters by scRNA-seq from PDAC organoids. **A.** Experimental design. Organoids were cultured from EUS-FNA samples, dissociated into single cells, splited into wells and labeled with well-specific barcodes. During the first round RT barcoded primers were added to the RNA, followed by ligation of a second and third barcode. The final cDNA library was sequenced on a NovaSeq platform. **B.** UMAP projections of combined single-cell profiles from the six patients. Each dot represents a single cell, and color refers different clusters. Number of cells in each cluster is indicated.

### Characterization of intra-organoid heterogeneity in 6 BDPCOs

Unsupervised clustering analysis was performed using the shared nearest neighbor modularity optimization based algorithm implemented in Seurat R package. This transcriptomic single cell analysis on BDPCO highlighted 4 different cell subtypes with different gene expression profiles identified as four cell clusters named C0 to C3, being C0, the biggest one, accounting for 5,341 cells (about 60% of the total). C1 accounted for 1,521 cells (17%), C2 for 1,174 cells (13%) and C3 for 898 cells (10%) (Figure 2B). The cells distributed between the four clusters showed very low overlapping with cluster C1 and were more distal in the spatial distribution generated by UMAP (uniform manifold approximation and projection). These diversified transcriptomic patterns displayed by different cell sub-groups indicate that human pancreatic tumor organoids maintain an important cell heterogeneity within the epithelial compartment of the tumor.

### Characterization of molecular markers in cell clusters

To deeply characterize this intra BDPCO heterogeneity, we performed a differential analysis of gene expression between all four clusters (Figure 3A, 3B and Table S1). Except cluster C0, all other clusters are characterized by a high expression of at least one specific molecular marker. In fact, C0 cluster is mainly characterized by low expression of the genes identified as markers in other clusters (Figure 3B). The most remarkable cluster is C1 which is not only the most distal but also the cluster best defined by many specific molecular markers expressed in the majority of their cells. These markers include the phosphodiesterase *PDE3A*, the helicase *HFM1, DLG2* a member of the membrane-associated guanylate kinase family and *SLCO5A1*, a solute carrier organic anion transporter. It is important to note that *PDE3A* and *HFM1* are two of the best basal-like markers identified by Nicolle et al. 2017. Cluster C1 was also characterized by a lower expression level in a particular set of genes compared to all other clusters. This low expression gene set includes *INO80D* a component of chromatin remodeling complex, *TERF2* a component of the telomere nucleoprotein complex and *CSMD1* which is a potential tumor suppressor as suggested by the fact that its expression in human breast cancer cells inhibited their aggressiveness, migration, adhesion and invasion (Escudero-Esparza et al., 2016) (Figure 3A, 3B and Table S1). Cluster C2 presented a high expression of *NEAT1* in 100% of the cells; *NEAT1* is a long non-coding RNA that regulates the transcription of genes involved in cancer progression (He et al., 2019; Zeng et al., 2020). The most specific markers of cluster C3 were ANKRD36, ANKRD36C, and ANKRD36B. These genes encode for ANRK proteins, three cell cycle-regulated kinases that appear to be involved in microtubule formation and/or stabilization at the spindle pole during chromosome segregation. Remarkably, supporting this fact, a recent study revealed that ANKRD36 is an oncogene whose expression was linked with poor prognosis in renal cell carcinoma (Yamada et al., 2018).

**Figure 3.**
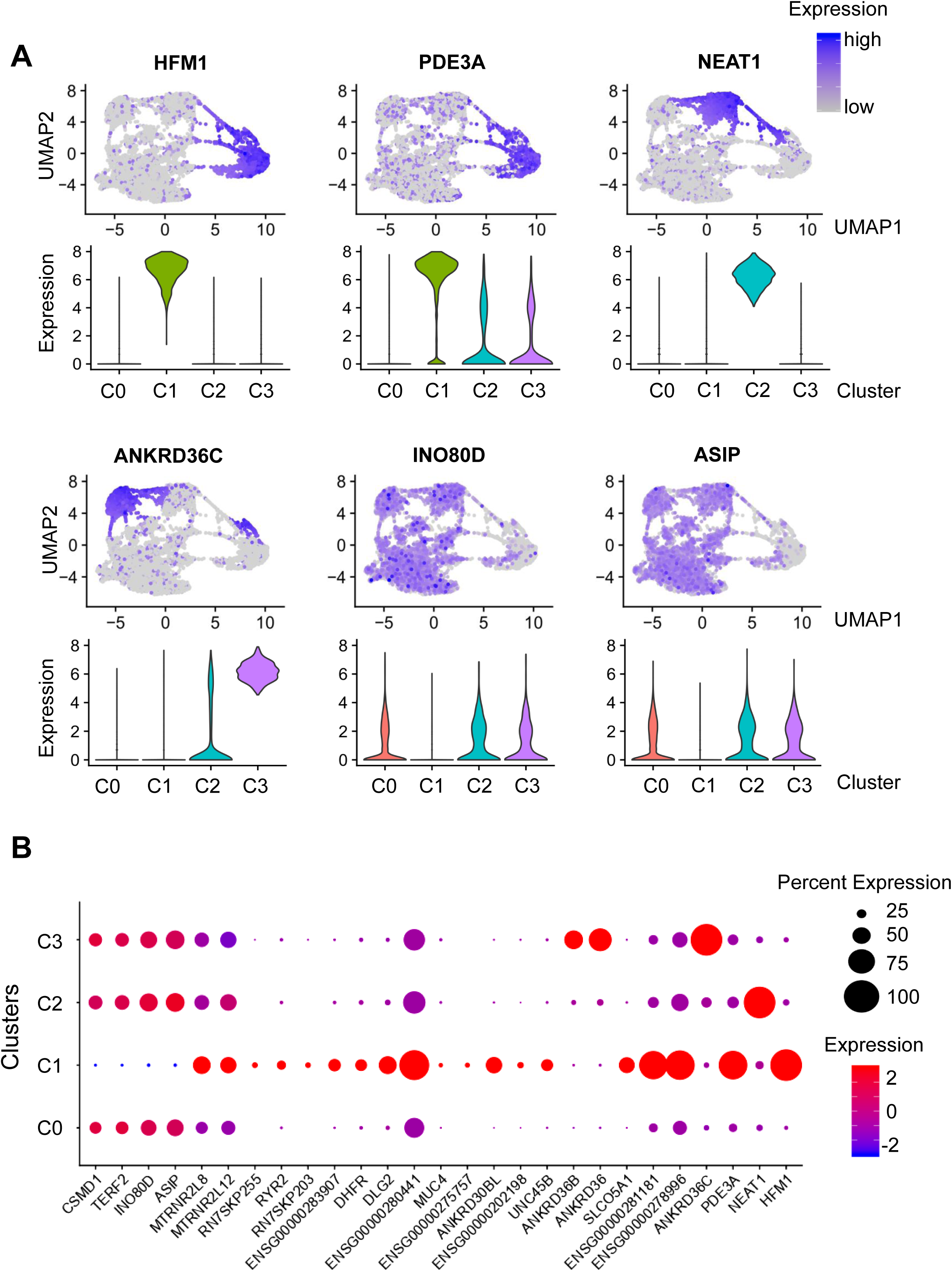
Characterization of cell clusters. **A.** Feature plot highlighting the expression of the gene markers for each cluster and the corresponding violin plots. **B.** Dot plot for the top markers of clusters. The dot size represents the percentage of cells expressing the marker and the colors indicate the expression level.

### Molecular components revealed the presence of basal-like cells in classical PDAC organoids

In order to annotate the single-cell clusters in accordance to their tumoral phenotype we extracted a basal-like and a classical PDAC component by performing an Independent Component Analysis (ICA) on the transcriptome dataset from the Nicolle et *al*. study (Nicolle et al., 2017). To validate the association of these two components to basal-like and classical phenotypes, we compared the correlation of the 1003 basal-like and 776 classical gene markers obtained from the Nicolle et *al*. work (Nicolle et al., 2017). As shown in Figure 4A by ICA analysis, the classical markers were significantly higher in the classical component compared to the basal-like (Student t-test p-value < 1e-16) and, as expected, the basal-like were significantly higher than classical in the basal-like component identified (t-test p-value < 1e-16). The projection of the average of the gene expression of clusters on these two components allowed the association of two molecular scores, the first corresponding to the classical and the second to the basal-like subtype. It is important to note that cluster C1 had the highest basal-like score while its classical score was the lowest among all other clusters. This supports the hypothesis that cluster C1 presents the most basal-like characteristics from all others. Cluster C2 presents a similar classical score when compared to C0 and C3 but was the cluster that had the lowest basal-like score (Figure 4B). Similar results were obtained at single-cell level by calculating the component scores of each individual cell (Figure 4C) confirming that cluster C1 contain basal-like cells that coexist with classical cells in all 6 BDPCOs. Cluster C1 contains 17% of all cells in our study which could represent the proportion of basal-like cells in these samples. The cluster C2 contains the most classical cells followed by C3 and then cluster C0. In terms of aggressiveness the single-cell clusters identified in this study could be ordered from the most to the least aggressive sub-group as follows: C1, C0, C3 and C2 respectively.

**Figure 4.**
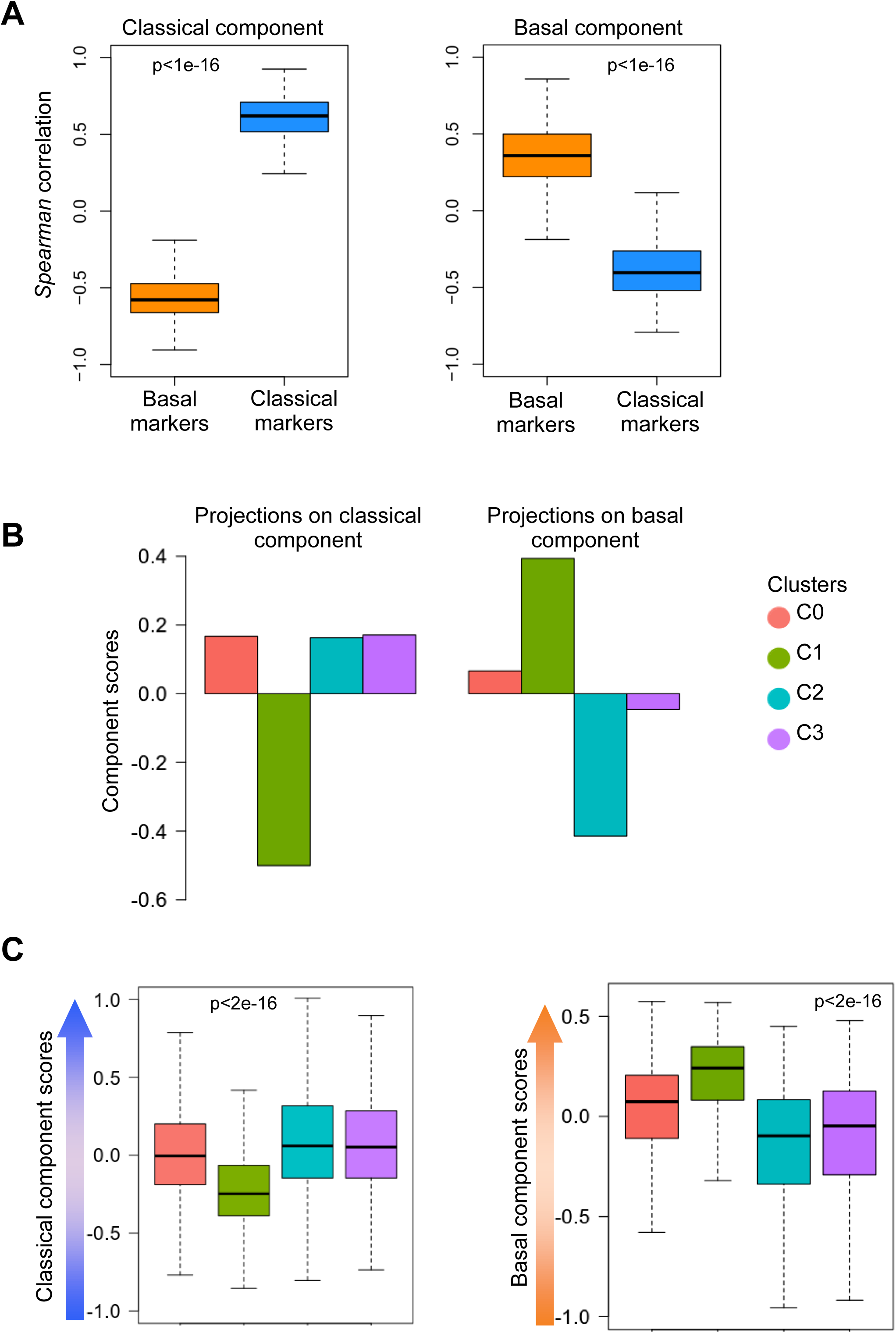
Identification and application of basal-like and classical components from transcriptomic to the single-cell data. **A.** Boxplots comparing basal-like and classical gene markers in the molecular components. **B.** Projections of average expression values of single-cell clusters on the components. **C.** Individual projections of each single-cell on the components. The boxplots of the single-cells projections were plotted by cluster.

### Analysis of pathway that characterize the different clusters

To characterize the biological profiles specifically associated with these different cell clusters, we performed pathway analysis using KEGG (Kyoto Encyclopedia of Genes and Genomes) (Figure 5A). Top enriched signaling pathways in cluster C0 were mainly the PI3K-Akt and Sphingolipid pathways. Studies showed that abnormal activation of the PI3K/AKT pathway promotes the proliferation of cancerous cells (Porta et al., 2014; Vasioukhin, 2012) Cluster C1 was characterized by a specific enrichment in many KEGG pathways including Adherents junction, Focal adhesion, Leukocyte trans endothelial migration, Glycolysis/Gluconeogenesis, Tight junction, etc. Most of these pathways are linked to each other as shown in the gene-function network (Figure 5B) and could indicate a high functional interaction of C1 cells with the extracellular matrix through which the cancerous cells have to migrate during the metastatic process (Maziveyi and Alahari, 2017; Vasioukhin, 2012). This analysis suggests that cluster C1 is highly invasive with a greater capacity to migrate and metastasize, this characteristic makes sense with the aggressive behavior of basal-like cells. Enriched pathways in cluster C2 were related to Mitophagy and AMPK signaling pathways among others. Previous studies have shown that activation of AMPK was an important regulator of mitochondrial homeostasis and that this activation initiates synthesis of new mitochondria to replace the damaged ones. In addition, activation of mitophagy pathways acts as key regulator of mitochondrial mass in cancerous cells as well as in homeostasis, bioenergetics, oncogene-driven metabolic reprogramming and cell apoptosis (Vara-Perez et al., 2019). Finally, cluster C3 was uniquely enriched in Neurotrophin signaling pathway. Recent studies have shown that the role of neurotrophins is not limited to neuronal tumors but also linked to nonneuronal tumors like thyroid, breast, lung, and prostate cancer (Tan et al., 2014).

**Figure 5.**
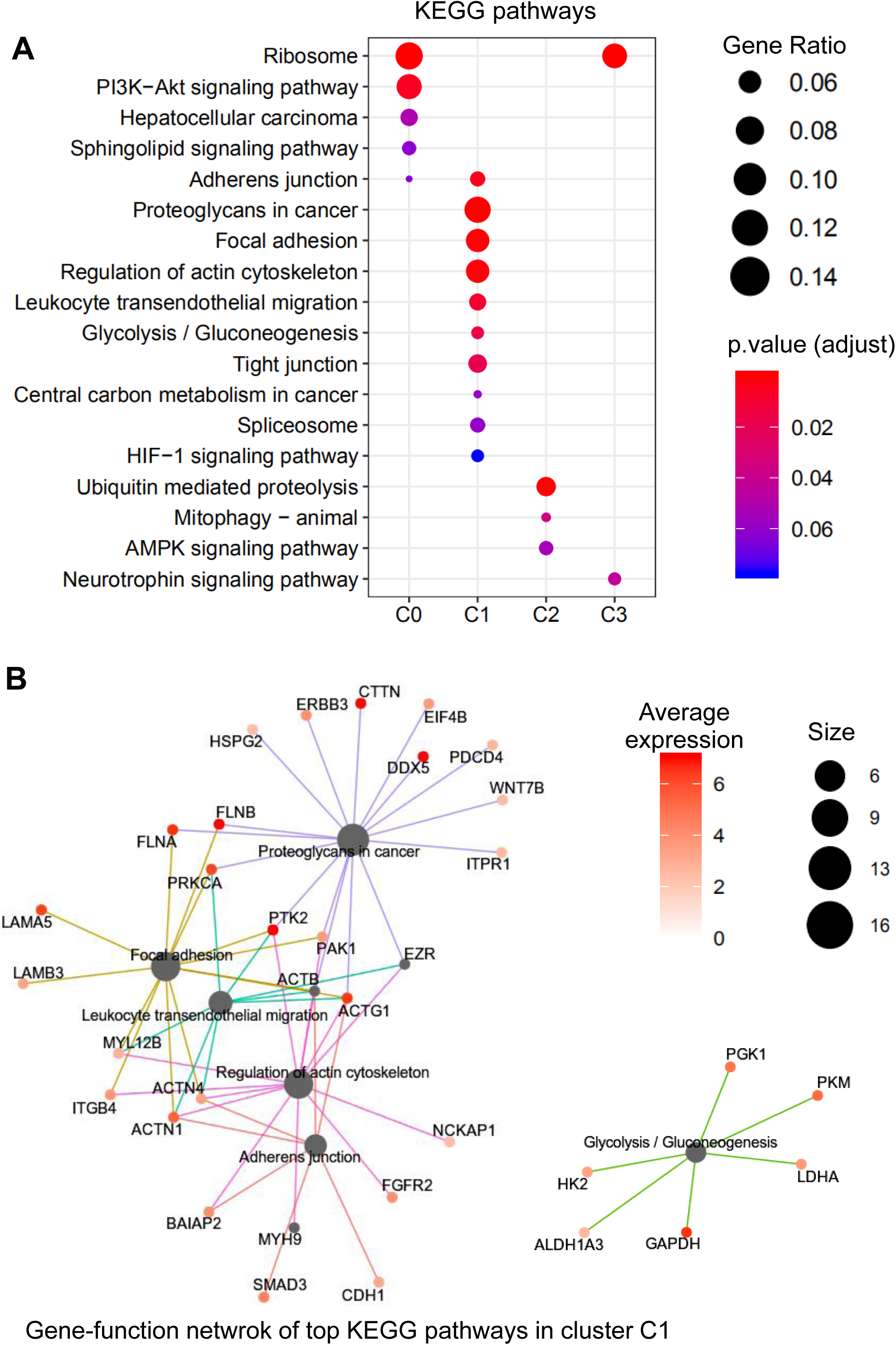
Pathway signatures of cell subtypes in organoids. **A.** Enrichment plot from KEGG pathway analysis comparing the single-cell clusters. **B.** Visualization of Gene-function network of top enriched pathways in cluster C1. The size is proportional to the number of genes associated to each pathway while the color indicates the average expression of genes in cluster C1.

### Pseudotime analysis of BDPCOs uncovers a differentiation trajectory

To unravel the putative developmental trajectory of cells during the tumorigenesis process we performed a single cell trajectory analysis using Monocle2 pseudotime trajectory (Qiu et al., 2017). The trajectory describes the virtual route through which the cells undergo changes during a defined biological process. Thus, the order of cells in the pseudotime trajectory depends on its particular state in this process. Our analysis resulted in a trajectory of multiple branches with cells from different clusters located in different places. This remote arrangement of clusters reflects the transcriptomic heterogeneity of the cells that compose each of them. As shown in Figure 6 and Figure S2, cells from different clusters were located in different branches of the trajectory especially sub-group C1 which presents the more distal location indicating a different state compared to others. Two main branches could be distinguished on the trajectory. The horizontal branches which contains principally the cells of cluster C1 and the vertical branches containing cells from other clusters. Clusters C2 and C3 were located in the bottom side of the horizontal branches while the cells of cluster C0 were located more on the top side. These observations lead us to hypothesize that the clusters could follow an organized temporal status during the tumorigenesis process as represented here by the pseudotime trajectory. Interestingly, the aggressiveness of cells (considering basal-like cells the most aggressive) seems to increase from the vertical-bottom (cluster C2 and C3) passing via vertical-top to the vertical left (cluster C0) to right branches (cluster C1) of the pseudotime trajectory (Figures 6, trajectory for each cluster and S2 trajectory for all clusters together).

**Figure 6.**
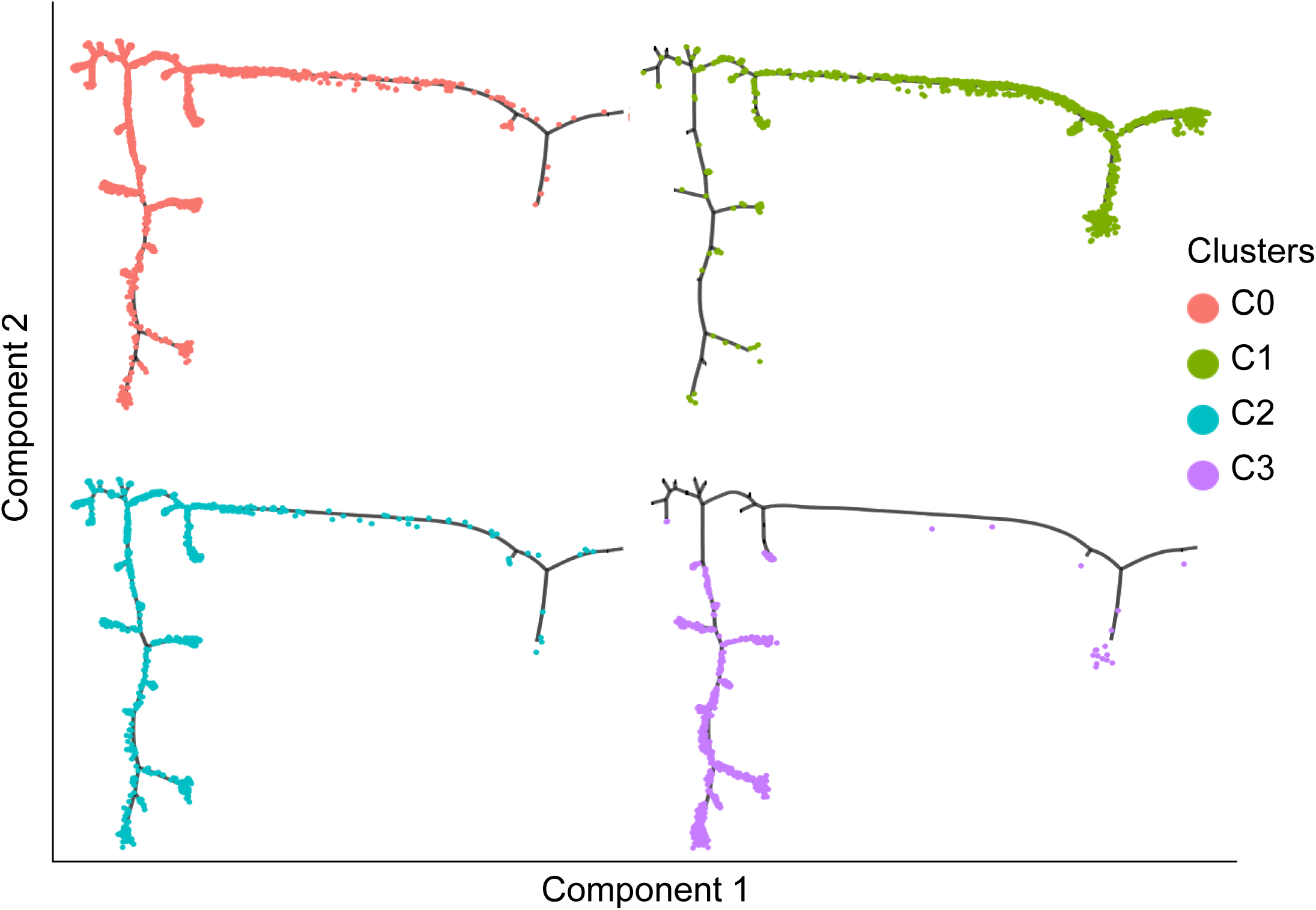
Pseudotime analysis of PDAC organoids. Pseudotime trajectory of single-cell transcriptomes simulating biological process in PDAC organoids. Each cluster of cells was plotted separately.

### Cell clusters in PDAC organoids are conserved across patients

By using combinatorial barcoding, we were able to determine the patient’s origin for each single cell in our study. In Figure 7, we split cells on the UMAP clustering according to patients and determined their proportions of each cluster. The four cell clusters found in this study were present in all six samples (Figure 7A and 7B) but in different proportions. In fact, and as an example, the proportion of cells that belonged to cluster C1, decreased from 20% in patient P2 to ∼12% in patient P1 (Table S3). To determine whether these different associations between patients and clusters are significant, we used *Pearson’s* Chi-squared test with simulated p-value based on 1000 replicates. We obtained a chi-square statistic of 53.2 and p-value < 1e-05 indicating that the cluster content statistically depends on the patient’s origin. Moreover, by using the *Pearson’s* residuals for which the absolute values indicate the contribution to the total Chi-square score above, we highlight the nature of dependence between each clusters and patients (Table S4). The results in Figure 7B showed the distinctive association between clusters across patients. For instance, cluster C1 had a strongest positive association with patient P2, then patient P4 and P6 respectively. However, this cluster (C1) had a repulsive relationship particularly with patients P1 then P3 and patient P5 at lower level. Other clusters were also associated distinctively to different patients such as C0 which was positively associated to P1 and negatively to P2 and P3 (Figure 7B). These results show that the intra-tumoral heterogeneity of PDAC cells is conserved across the patients but the content in the different clone vary between patients highlighting the heterogeneity between PDAC patients.

**Figure 7.**
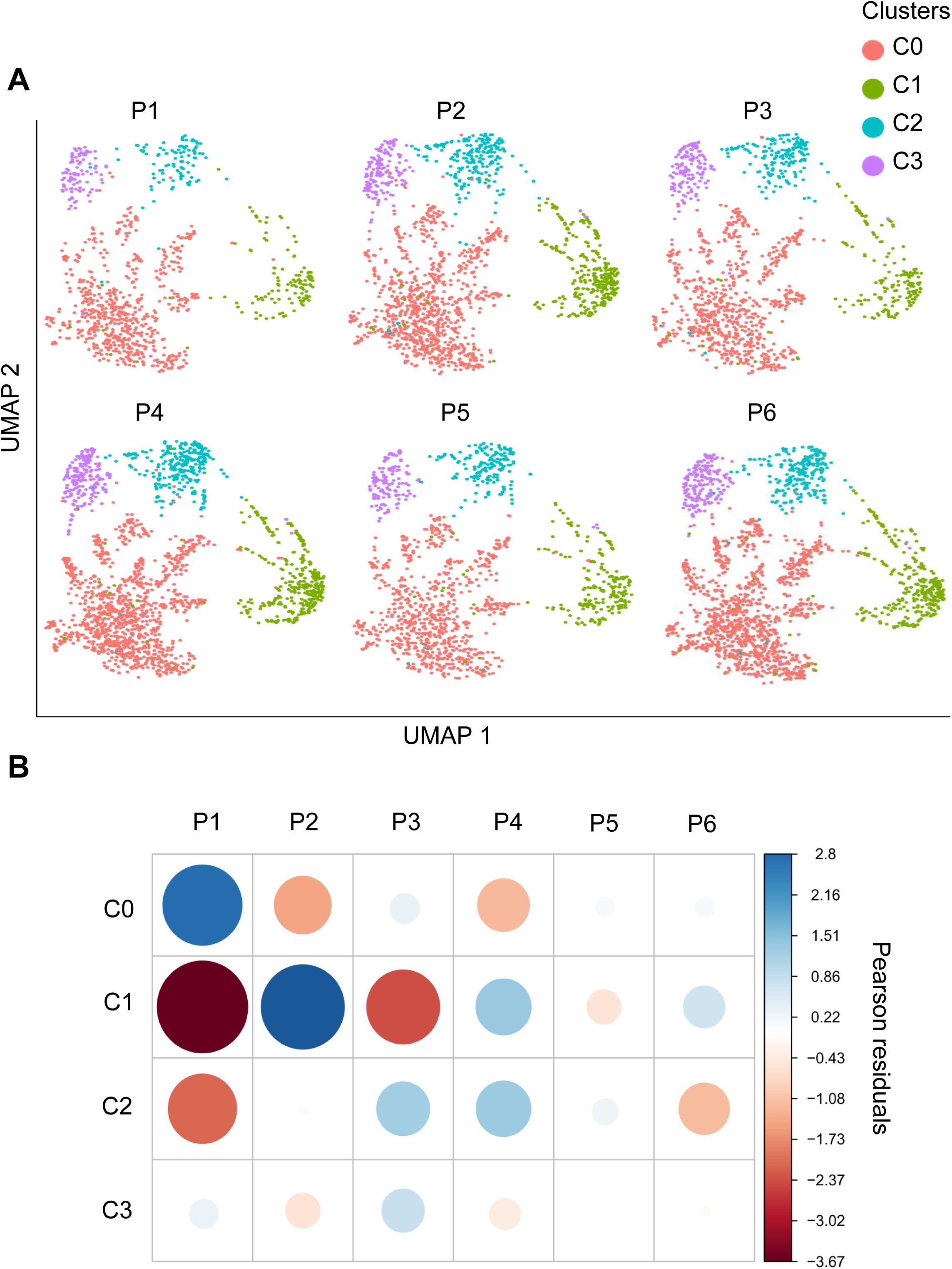
Intra-tumoral heterogeneity is conserved across the patients with different proportions. **A.** UMAP projections of combined single-cell clusters separately for each patient. **B.** Correlogram plot of *Pearson’s* residuals from Chi-squared test between the cluster and patients. The color of circles indicates the nature of relationship between the patients and clusters while the absolute value indicates the global contribution to Chi-square score.

## Discussion

One of the main difficulties in finding an effective treatment for PDAC is its heterogeneity. PDAC is currently stratified into two main different phenotypes: basal-like and classical, based on molecular subtyping by gene expression profiling (Moffitt et al., 2015; Nicolle et al., 2017). However, this practical classification does not take into account the heterogeneity that exists within each tumor and many sources of evidence indicate that mixed tumors (containing basal-like and classical cells) could exist. Characterizing this intra-tumor heterogeneity is essential for really understanding PDAC evolution and to envision new insights that will conduce to more personalized and efficient therapies. Recently, it became possible to investigate intra-tumor heterogeneity at a single-cell resolution identifying different cell types in PDAC and opening a way to study distinct functions of cancer-associated fibroblasts subtypes in PDAC immunity and progression (Elyada et al., 2019). In fact, human primary tumor from surgery, pancreatic biopsies obtained by EUS-FNA and/or xenografts contains many different types of cells other than epithelial (fibroblasts, immune infiltrate, blood cells, etc) which can significantly impact the study of the differences between tumor cells in single-cell experiments. However, the organoid is an excellent model for in depth analysis of pure epithelial tumor cells allowing the study of intrinsic epithelial heterogeneity in a single pancreatic tumor (Brazovskaja et al., 2019). From a methodological standpoint, the study presented here represents a proof of concept that SPLIT-seq technique on BDPCO can be used to deeply characterize tumor heterogeneity in the epithelial cell compartment of PDAC. We choose this approach because it presents many advantages for studying PDAC heterogeneity on samples directly obtained from patients. It includes the possibility of studying a high number of cells in a single experiment obtaining up to 884,736 unique barcode combinations after 3 ligations, profiling several samples in parallel thus reducing the batch effect, and better preservation of the transcriptomes by reducing the steps required before cell fixation.

In our study, we characterized six consecutive PDAC tumors from patients to analyze their intra-tumor heterogeneity. The 6 tumors used in our study were of classical subtype as suggested by our histologic, immunofluorescence and transcriptomic analyses. It is important to note that we detected some cells expressing VIM, as a basal-like marker, in all these organoids, as well as in the organoids-derived PDXs, after at least two consecutive passages in mice, indicating that the intra-tumor heterogeneity in PDAC is frequent if not systematic. At the scRNA analysis we identified four cell clusters or subpopulations, using a well-defined bioinformatics set-up, in all six patients analyzed. Remarkably, these clusters are recurrently present in the PDAC tumors although in different proportions, suggesting that the aggressiveness of the PDAC could be controlled, at least in part, by the presence of the most aggressive subpopulation, probably the C1. In addition, another source of the intra-tumor heterogeneity of the epithelial cells, which was not considered here, may originate from the local differences of the tumor.

In this study we intended to highlight for the first time the intrinsic heterogeneity within the epithelial cancerous cell compartment of 6 classical PDAC patients. We observed that the cluster C0 contains most of the cells studied which share a common transcriptomic profile.

Other clusters were characterized by the expression of specific transcriptomic markers expressed in the majority of cells. In many cases, these markers have been previously related to tumor specific biology. For instance, *NEAT1* (marker of cluster C2) has shown to be up-regulated in cancer and plays a role in most types of solid tumors by regulating tumor suppressive microRNAs (Yu et al., 2017), *NEAT1* has been suggested as a marker of poor prognosis in colorectal cancer (Li et al., 2015) and glioma (He et al., 2019). Cells of cluster C1 were also characterized by several specific molecular markers such as for example *PDE3A* that encodes a protein which controls degradation of cyclic AMP (cAMP) and GMP (cGMP) (Beavo, 1995) that regulates various physiologic processes including adherents junction, glycolysis/gluconeogenesis and leukocyte transendothelial migration pathways. This high expression suggests that C1 cells have high metabolic and migration activities, which could correspond to highly aggressiveness cells with strong metastatic potential, supporting the basal-like phenotype of this cluster. Altogether, the diversified transcriptomic patterns displayed by different clusters indicate that PDAC organoids maintain important cell heterogeneity and that the presence of basal-like cells within all PDAC tumors studied here brings new insights into the intra-tumoral heterogeneity in PDAC cancer.

In a recent work, Peng et al. (Peng et al., 2019) ports a study on single cell transcriptome analysis of a total of 57,530 cells from 24 primary PDAC tumors and 11 control pancreases. They found that PDAC tumor mass is highly heterogeneous and composed of diverse malignant and stromal cell types as expected. In addition, they report that malignant ductal subtype could be distinguished by featured gene expression profile and was observed to contain highly proliferative and migratory subpopulations. They suggest that these cell subtypes could correspond to the basal-like, which represents around 6.30% of the ductal in their samples, and classical subtype which represent 26.95% of the cells, however their protocol setup was directed mainly to describe the cellular composition of the PDAC.

In summary, scRNA-seq analysis performed on 6 consecutives PDAC as organoids allowed us the identification of four main cell clusters present in different proportions in all tumors. Clusters show a specific gene expression profile associated with specific biological characteristics and molecular markers. Although these tumors were classified as Classical when analyzed in bulk, one of the clusters present in all of the patients, corresponded to a basal-like phenotype. These results depict the unanticipated high heterogeneity of pancreatic cancers and demonstrated that basal-like cells with a highly aggressive phenotype are more widespread than expected. We conclude that Basal-like and Classical cells coexist in PDAC.

## Acknowledgments

This work was supported by INCa (Grants number 2018-078 and 2018-079), Canceropole PACA, Amidex Foundation, Fondation de France, La Ligue Contre le Cancer and INSERM. Authors wish to thanks Christopher Pin for critically reading the manuscript.

## Author Contributions

The study was designed by N. D., R. N. and J. I. The experiments were conducted by N. J., J. R., M. B. and O. G. Data was analyzed by A. E. and R. N. The manuscript was written by N. J., A. E., J. I. and N. D. All authors read and approved the final manuscript.

## MATERIALS AND METHODS

### Samples

Patients were included under the Paoli Calmettes Institute clinical trial NCT01692873 (https://clinicaltrials.gov/show/NCT01692873). Consent forms of informed patients were collected and registered in a central database.

Primary PDAC-derived organoids were obtained from consecutive patients with unresectable tumors by endoscopic fine-needle aspiration (EUS-FNA). Biopsies were slightly digested with the Tumor Dissociation Kit, human (Miltenyi Biotec) at 37°C for 5 min, then incubated with Red Blood Cell Lysis Buffer (Roche), and washed two times with PBS. Samples were placed into 12-well plates coated with 150 µl GFR matrigel (Corning) and cultured with pancreatic organoid feeding media (advanced DMEM/F12 supplemented with 10 mM HEPES; Thermo-Fisher); 1x Glutamax (Thermo-Fisher); penicillin/streptomycin (Thermo-Fisher); 100 ng/ml animal-free recombinant human FGF10 (Peprotech); 50 ng/ml animal-free recombinant human EGF (Peprotech); 100 ng/ml recombinant human Noggin (Biotechne); Wnt3a-conditioned medium (30% v/v); RSPO1-conditioned medium (10% v/v); 10 nM human Gastrin 1 (Sigma Aldrich) 10 mM Nicotinamide (Sigma Aldrich); 1.25 mM N acetylcysteine (Sigma Aldrich); 1x B27 (Invitrogen); 500 nM A83-01 (Tocris); 10.5 µM Y27632 (Tocris). The plates were incubated at 37°C in a 5% CO_2_ incubator, and the media changed every 3 to 4 days.

### Immunohistochemistry

Organoids and PDX were embedded, section and stained for H&E and/or histology. Immunofluorescent staining with COL-IV and ZO-1 antibodies was performed using anti-collagen IV rabbit polyclonal antibody (Abcam, ref ab6586), anti-ZO1 monoclonal antibody (ThermoFisher, ref Z01-1A12) and Anti-Vimentin monoclonal antibody (Sigma, ref. V6389) following standard methods.

### Single-cell transcriptomics

For single-cell transcriptomic, we performed a modified SPLIT-seq protocol similar to the already described by Rosenberg et *al*. (Rosenberg et al., 2018). Briefly, single-cell suspensions from organoids were fixed with 1% formaldehyde solution in PBS and stored at −80°C immediately after dissociation. All samples were thawed in ice and permeabilized with 0.2% Triton X-100. Samples were divided into a 48-well plate. Retro-transcription was performed with well-specific barcoded oligo(dT) and hexamer primers. Cells were pooled and split twice to 96-well plates for two successive ligation steps where a second and a third well-specific barcodes were added to the cDNA. For library preparation, two rounds of SPRI size selection were performed after cDNA amplification and the tagmentation steps. Illumina amplicons were generated from 1 ng instead of 600 pg of cDNA and sequenced using the Illumina Novaseq platform.

### scRNA-seq data processing

Raw sequencing data were processed using the zUMIs pipeline (Parekh et al., 2018) which consisted mainly in extracting barcodes, filtering cells, mapping using human genome GRCh38.96 and generating read count tables. Downstream analyses on gene-by-cell count matrix were performed with the R package Seurat, version 3 (Butler et al., 2018). These consist mainly of normalization, dimensional reduction (including PCA and UMAP algorithm) and cell classification using a shared nearest neighbor (SNN) modularity optimization based clustering algorithm. Markers for each cluster were identified based on differential gene expression using the Seurat’s function FindMarkers. Pseudotime analysis was performed using the R package Monocle2 trajectory (Qiu et al., 2017). Enrichment analyses including Gene Ontology and KEGG Pathway were performed using R packages such as ClusterProfiler (Yu et al., 2017).

### Identification of basal-like and classical components

Transcriptomic dataset was pre-processed and normalized using only 50% most variable genes. Thus, data were thus sample-wise zero-centered and scaled. Independent component analysis was performed with JADE (joint approximate diagonalization of eigenmatrices) algorithm. Two components were retrieved using biologically relevant composition. The annotation of the components was performed using external data including PDX samples and lists of basal-like and classical markers. The cluster projections and gene correlation to the components was performed using custom R scripts and basic functions of R language.

---

**Supplementary Figure S1.**
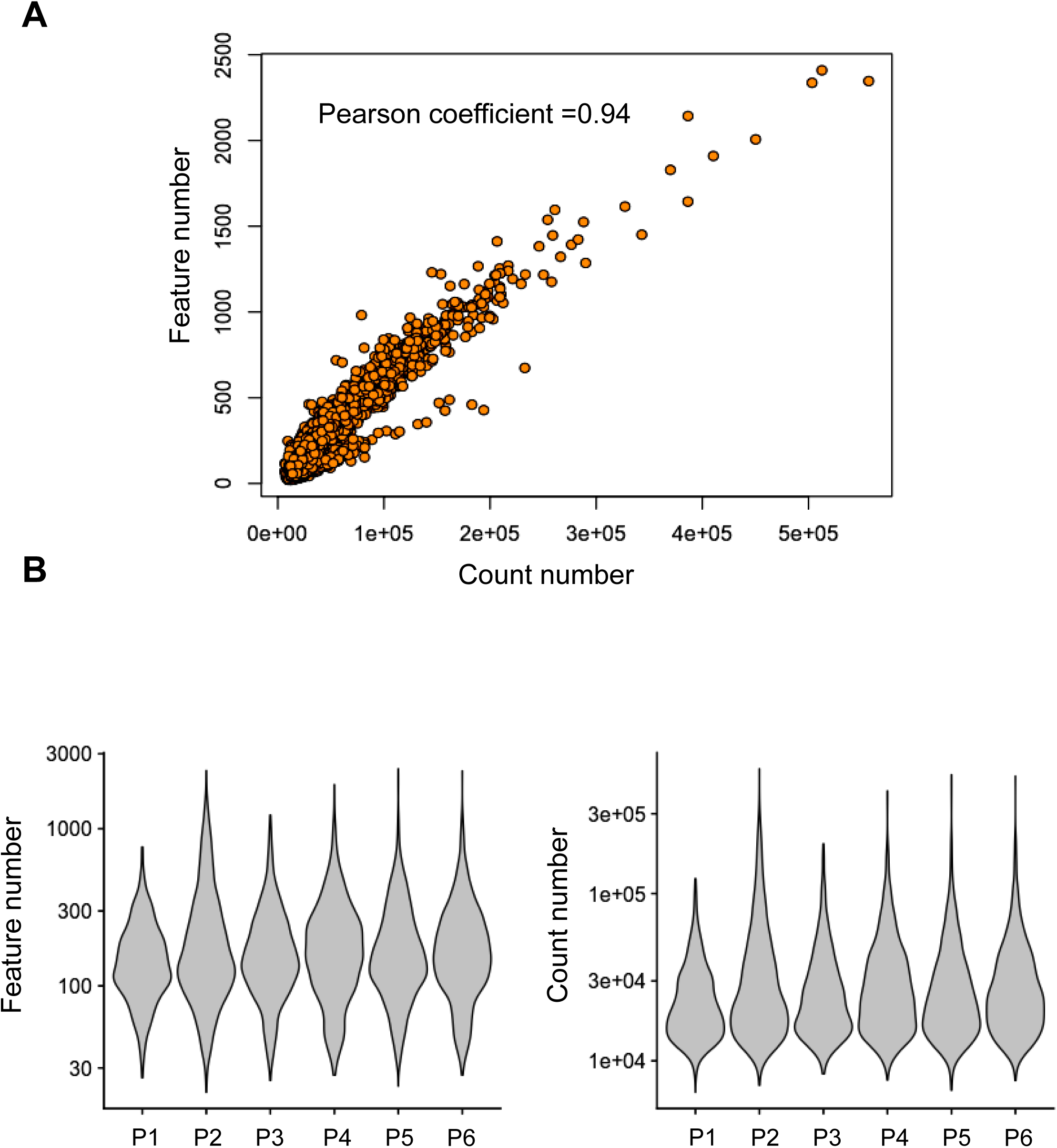
Performance of scRNA-seq by combinational indexing. **A.** Scatter plot showing the correlation between read counts and genes (Pearson coefficient =0.94). **B.** Violin plots of detected genes and number of reads across patients.

**Supplementary Figure S2.**
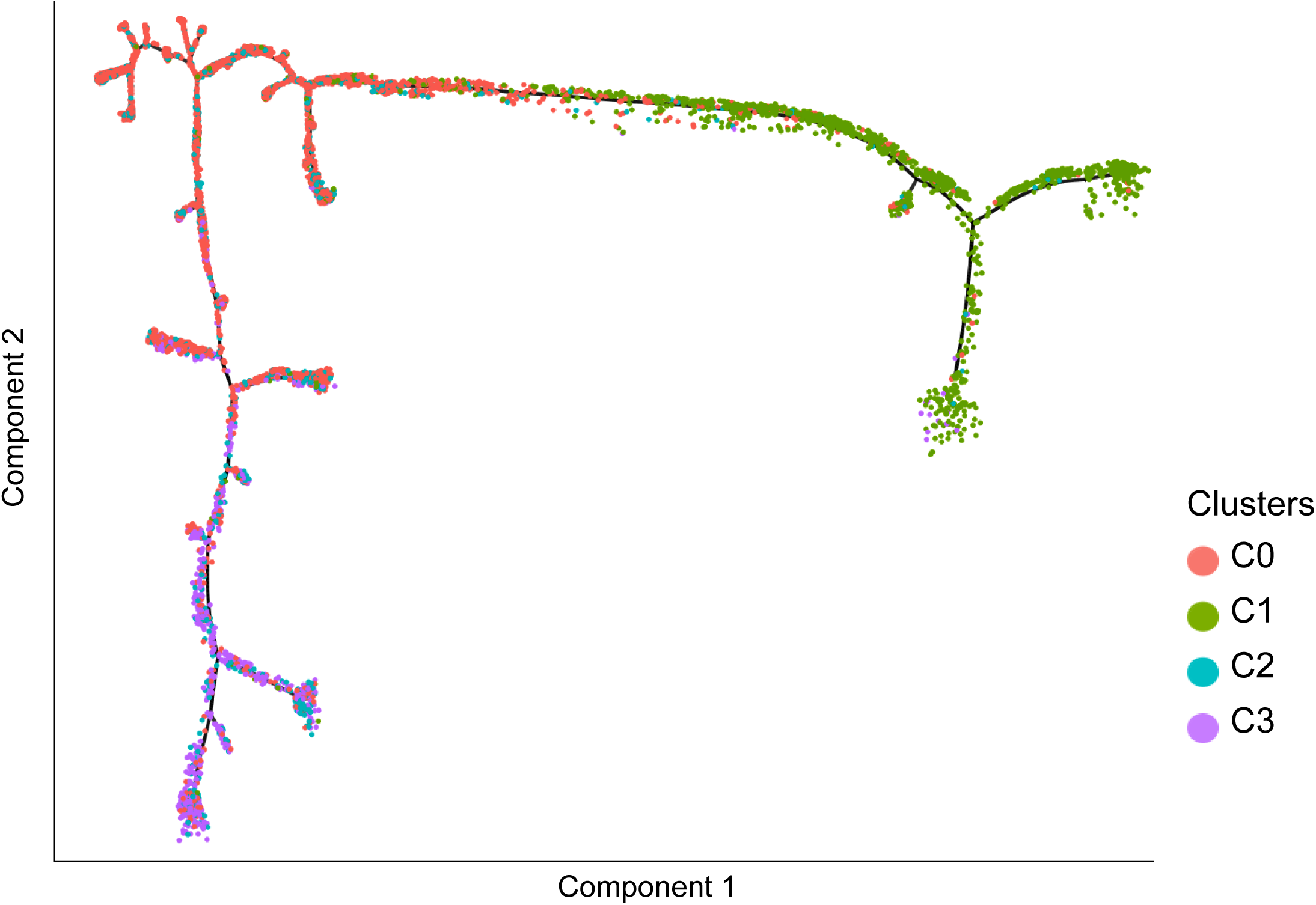
Pseudotime trajectory. Pseudotime trajectory analysis indicating the state of all cells from the four clusters together on the same trajectory.

